# A systematic benchmark of Nanopore long read RNA sequencing for transcript level analysis in human cell lines

**DOI:** 10.1101/2021.04.21.440736

**Authors:** Ying Chen, Nadia M. Davidson, Yuk Kei Wan, Harshil Patel, Fei Yao, Hwee Meng Low, Christopher Hendra, Laura Watten, Andre Sim, Chelsea Sawyer, Viktoriia Iakovleva, Puay Leng Lee, Lixia Xin, Hui En Vanessa Ng, Jia Min Loo, Xuewen Ong, Hui Qi Amanda Ng, Jiaxu Wang, Wei Qian Casslynn Koh, Suk Yeah Polly Poon, Dominik Stanojevic, Hoang-Dai Tran, Kok Hao Edwin Lim, Shen Yon Toh, Philip Andrew Ewels, Huck-Hui Ng, N.Gopalakrishna Iyer, Alexandre Thiery, Wee Joo Chng, Leilei Chen, Ramanuj DasGupta, Mile Sikic, Yun-Shen Chan, Boon Ooi Patrick Tan, Yue Wan, Wai Leong Tam, Qiang Yu, Chiea Chuan Khor, Torsten Wüstefeld, Ploy N. Pratanwanich, Michael I. Love, Wee Siong Sho Goh, Sarah B. Ng, Alicia Oshlack, Jonathan Göke, SG-NEx consortium

**Author notes:** nf-core/nanoseq is a streamlined, community curated pipeline for Nanopore Sequencing data processing and analysis: https://nf-co.re/nanoseq The SG-NEx data is available: https://github.com/GoekeLab/sg-nex-data.

## Abstract

The human genome contains more than 200,000 gene isoforms. However, different isoforms can be highly similar, and with an average length of 1.5kb remain difficult to study with short read sequencing. To systematically evaluate the ability to study the transcriptome at a resolution of individual isoforms we profiled 5 human cell lines with short read cDNA sequencing and Nanopore long read direct RNA, amplification-free direct cDNA, PCR-cDNA sequencing. The long read protocols showed a high level of consistency, with amplification-free RNA and cDNA sequencing being most similar. While short and long reads generated comparable gene expression estimates, they differed substantially for individual isoforms. We find that increased read length improves read-to-transcript assignment, identifies interactions between alternative promoters and splicing, enables the discovery of novel transcripts from repetitive regions, facilitates the quantification of full-length fusion isoforms and enables the simultaneous profiling of m6A RNA modifications when RNA is sequenced directly. Our study demonstrates the advantage of long read RNA sequencing and provides a comprehensive resource that will enable the development and benchmarking of computational methods for profiling complex transcriptional events at isoform-level resolution.

## Introduction

Each cell type and tissue has a specific gene expression profile, and differences in the set of transcribed genes provide high dimensional insights into cell identities and alterations in diseases. In the human genome, protein coding genes can generate on average 7 different gene isoforms ^1^. Alternative promoters, exon skipping, intron retention, 3’ end sites and polyadenylation all contribute to this diversity, enabling a single gene to generate a large number of distinct transcripts. Differences in gene isoform expression have been observed and are found to be highly informative in cancer and other diseases even when the overall gene expression levels are stable ^2–7^.

The comprehensive profiling and low cost of short read RNA-seq have made it one of the most widely used technologies to study molecular properties of cells and tissues ^8^. The majority of short read RNA-Seq data is based on PCR amplified sequencing of cDNA, which introduces biases that lead to different exons being sequenced at different coverage levels ^9^. While short read data generates robust estimates for gene expression, the presence of overlapping annotations and systematic biases limits the ability to uniquely assign reads to individual gene isoforms ^10,11^. To deal with the increased uncertainty, approaches have been developed that focus on specific splice-junction or exon usage ^2,6,12,13^, however, more complex transcriptional events that involve multiple exons are often not fully captured ^14–19^.

Transcriptome profiling using Nanopore long read sequencing promises to overcome some of the main limitations of current short read RNA-Seq protocols ^20–24^. While nanopore sequencing is prone to a higher error rate ^25^, the sequencing platform is one of the most widely available, potentially enabling long read RNA sequencing at a cost per gigabase comparable with current short read technologies. The Oxford Nanopore sequencing platform performs long read RNA sequencing with 3 different protocols. PCR-amplified cDNA sequencing (PCR-cDNA) requires the least amount of input RNA and generates the highest throughput. When sufficient RNA is available, the PCR step can be omitted using the direct cDNA protocol. The direct RNA-Seq protocol enables sequencing of native RNA, thereby avoiding the reverse transcription and amplification steps as well as providing information about possible RNA modifications ^26^. While several long read RNA-Seq data sets have been described they are either low throughput ^21,23,24,26^, covering single conditions ^24^, or individual protocols ^26^, limiting the ability to comprehensively compare and evaluate the different RNA-Seq protocols.

Here we present the results from the Singapore Nanopore Expression Project (SG-NEx), a comprehensive benchmark data set and systematic comparison of the different Nanopore RNA-Sequencing protocols. The SG-NEx data consists of 5 human cell lines that were sequenced in multiple replicates using direct RNA, amplification-free direct cDNA, and PCR-amplified cDNA long-read protocols, as well as short read RNA-seq. The data includes spike-in RNAs with known concentrations ^27^. We further provide a community-curated nf-core pipeline that simplifies data processing, methods evaluation and discovery. Our results indicate that long read sequencing improves transcript quantification and facilitates novel analysis of full length isoforms, RNA modifications, and fusion genes that are impossible with current short read technology. Our study provides a data-driven guide for experimental design of transcriptomics experiments, and describes a resource that will be invaluable for benchmarking and development of computational methods for transcript discovery and quantification, differential expression analysis, fusion gene detection and identification of RNA modifications from long read RNA-Seq data.

## Results

### (1) Deep transcriptome profiling of human cell lines using Nanopore RNA-Sequencing

To provide a comprehensive benchmark data set for Nanopore RNA-Seq data, we have deeply profiled one common cell line for colon cancer (HCT116), liver cancer (HepG2), lung cancer (A549), breast cancer (MCF7), and Leukemia (K562) with at least 3 high quality replicates using the direct RNA protocol (direct RNA), the amplification free cDNA protocol (direct cDNA), the PCR cDNA protocol (cDNA), and short read Illumina cDNA Sequencing (Figure 1a). For a subset of sequencing runs, we included sequin spike-in RNAs with known concentrations that enable the evaluation of transcript discovery and quantification ^27^. In total, we have sequenced 88 libraries, with an average sequencing depth of 34 million long reads per cell line (Figure 1b,c, Supplementary Table 1).

**Figure 1.**
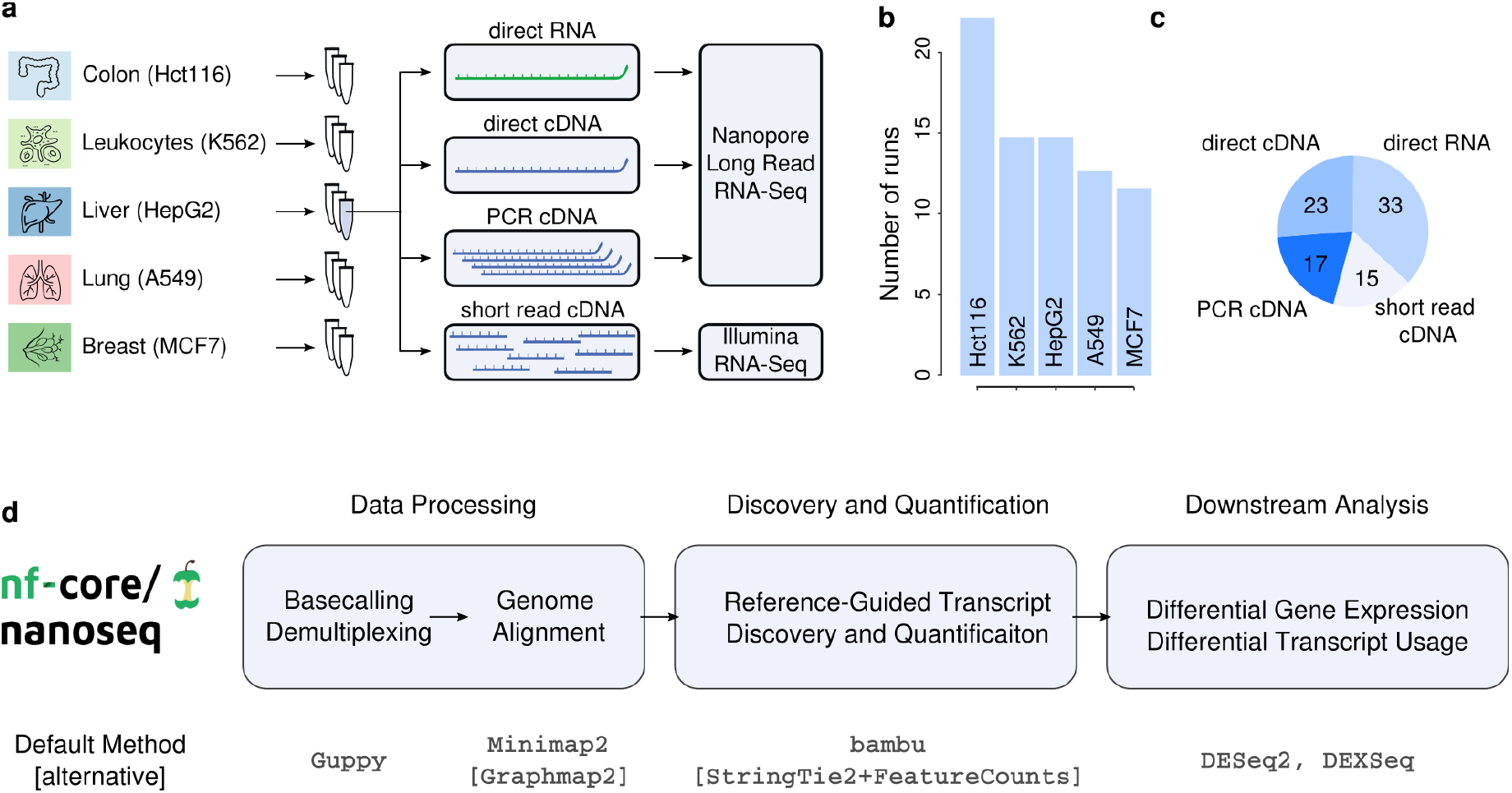
Overview of the Singapore Nanopore Expression (SG-NEx) datasets and processing pipeline. **(a)** 5 cancer cell lines were sequenced with multiple replicates using 4 different RNA-Seq protocols. **(b)**Shown is the number of sequencing runs generated for each SG-NEx cell line. **(c)** Shown is the number of sequencing runs for each of the sequencing technologies. **(d)** Illustration of the nf-core nextflow pipeline (nanoseq) for streamlined processing of Nanopore long read RNA-Seq data.

### (2) nf-core/nanoseq: a community curated pipeline for Nanopore data processing

To facilitate the streamlined processing and analysis of long read RNA-Seq data, we developed the nanoseq pipeline (Figure 1d). Nanoseq consists of 3 different modules: Basecalling/Alignment, Transcript Discovery/Quantification, and Differential Expression/Differential Transcript Usage (Figure 1d). Each module provides the option to use different existing methods that can be seamlessly integrated with the other modules. This design ensures consistency in processing RNA-Seq data, enabling the benchmarking of different sequencing protocols and computational methods. The pipeline is dynamically tested on a full size data set, it allows data processing through Docker and Singularity, and can be executed on the cloud. Nanoseq is implemented in the nextflow language ^28^ and maintained as a community curated pipeline on nf-core ^29^ (https://nf-co.re/nanoseq).

### (3) A benchmark of Nanopore long read RNA-Seq protocols

Using the nanoseq pipeline we processed all SG-NEx Nanopore RNA-Seq data to obtain a curated set of transcript annotations and quantification estimates that allow us to compare the different Nanopore long read RNA-Seq protocols in terms of throughput, read length, transcript coverage, and potential sequencing biases. Globally, we found that gene expression estimates are consistent across all sequencing protocols and platforms (Figure 2a). Biological variation dominates gene expression, whereas technical variation due to the sequencing platform is minimal (Supplementary Figure 1a). Among the long read RNA-Seq protocols, PCR amplified cDNA sequencing consistently generated the highest throughput (Figure 2b). However, not all genes appear to be equally amplified in the PCR step. The top 1,000 most highly expressed genes across all samples represented 69% of all reads in the PCR-cDNA data, compared to 62% for direct RNA and 59% for direct cDNA data (Figure 2c, Supplementary Figure 1b). This suggests that the increased sequencing depth through PCR amplification comes at the cost of lower transcriptional diversity, with highly expressed genes being over-represented compared to the PCR-free protocols.

**Figure 2.**
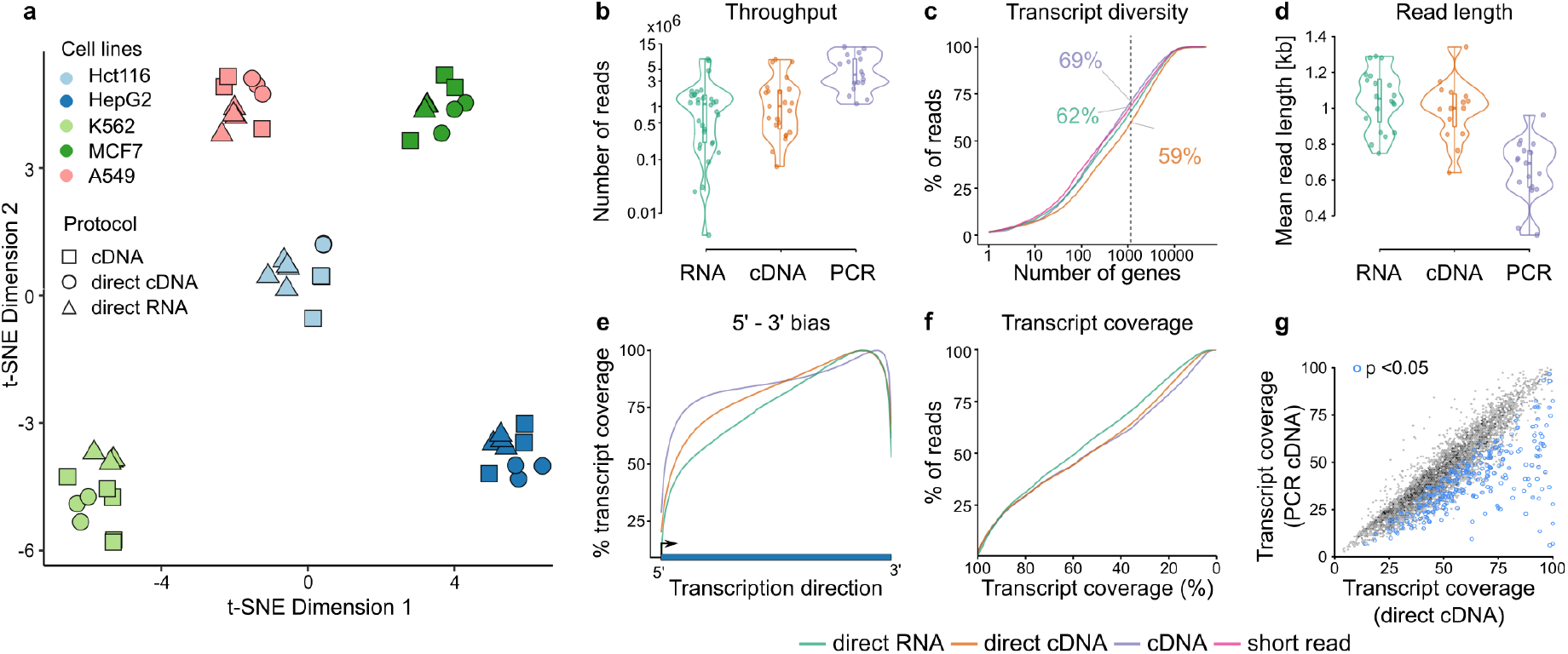
Comparison of Nanopore RNA-Seq protocols. **(a)** Shown is the t-SNE plot of gene expression across all the samples generated using cDNA, direct cDNA and direct RNA protocols from the 5 cancer cell lines. **(b)** Violin plot showing the sequencing throughput of RNA (direct RNA), cDNA (direct cDNA) and PCR (cDNA) protocols. **(c)** Shown is the transcription diversity depicted by the fraction of reads attributed to the number of genes ranked by expression levels from highest to lowest for the three protocols. The dotted line represents the top 1000 expressed genes and numbers in color indicating the fraction of reads accounted for them. **(d)** Violin plot plot showing the average read length per sample of RNA (direct RNA), cDNA (direct cDNA) and PCR (cDNA) protocols. **(e)** Shown is the average coverage along the normalized transcript length for RNA (direct RNA), cDNA (direct cDNA) and PCR (cDNA) protocols. **(f)** Cumulative distribution plots showing the proportion of reads that cover at least the given fraction of transcripts, for RNA (direct RNA), cDNA (direct cDNA) and PCR (cDNA) protocols. **(g)** Shown is the transcript coverage of reads generated using the direct cDNA and the PCR cDNA protocol. Blue points indicate genes with significantly reduced coverage in the PCR cDNA protocol.

Among the Nanopore RNA-Seq protocols, direct RNA-Seq generated the highest average read length (Figure 2d). The direct RNA-Seq protocol starts the sequencing process at the polyA tail and we observe an increased coverage at the 3’ end compared to the cDNA protocols as expected (Figure 2e). However, despite this 3’ sequencing bias, the direct RNA-Seq protocol still shows the highest average transcript coverage, partially due to increased read length (Figure 2f). Interestingly, a comparison of the average base coverage for each gene suggests that certain genes are incompletely sequenced in the PCR cDNA protocol (Figure 2g, Supplementary Figure 1c). In particular, a set of 362 genes showed a significant decrease in bases covered that suggests a bias for the cDNA-PCR protocol that might affect the accurate quantification of their transcripts (Supplementary Table 2). The reduced coverage appears to be linked to the presence of low complexity sequences and amplification of short fragments from the 3’ end among others (Supplementary Figure 1d). Globally, we find that all 3 protocols provide comparable gene and transcript expression estimates, with the direct RNA and direct cDNA protocols being most similar (R=0.937, Supplementary Figure 1e), possibly reflecting a more unbiased representation of the transcriptome.

### (4) Short and long read RNA-Seq provide comparable estimates for gene expression analysis

The ability to robustly estimate gene expression and identify differentially expressed genes has made short read RNA-Seq data widely popular. To compare the ability of long and short read RNA-Seq data to quantify gene expression, we first analysed estimates for spike-in data. Across the spike-in genes, estimates from long read RNA-Seq data showed lower estimation error and a higher correlation with the expected concentrations (R = 0.97 vs 0.93 Figure 3a,b, Supplementary Figure 2a,b). Nevertheless, gene expression estimates from short and long read RNA-Seq were still highly consistent both for spike in RNAs (R = 0.93, Figure 3c) and non-spike in RNAs (R = 0.92, Figure 3d, Supplementary Figure 2c). On RNA from human cell lines we only observed systematic differences of expression estimates for pseudogenes which have higher read counts with long read data. This could be due to improved resolution at repetitive regions or higher error rates of long reads (Figure 3e, Supplementary Figure 2d). The high consistency between short and long read data is also reflected in differential expression estimates when different cell lines are compared (Figure 3f, Supplementary Figure 2e). While the set of genes at specific p-value thresholds differed, the ranking of differentially expressed genes was largely preserved (Supplementary Figure 2f). Differential expression results were also similar when long and short read data were jointly analysed, indicating that Nanopore RNA-Seq data can be integrated with existing short read RNA-Seq data (Figure 3g, Supplementary Figure 2g).

**Figure 3.**
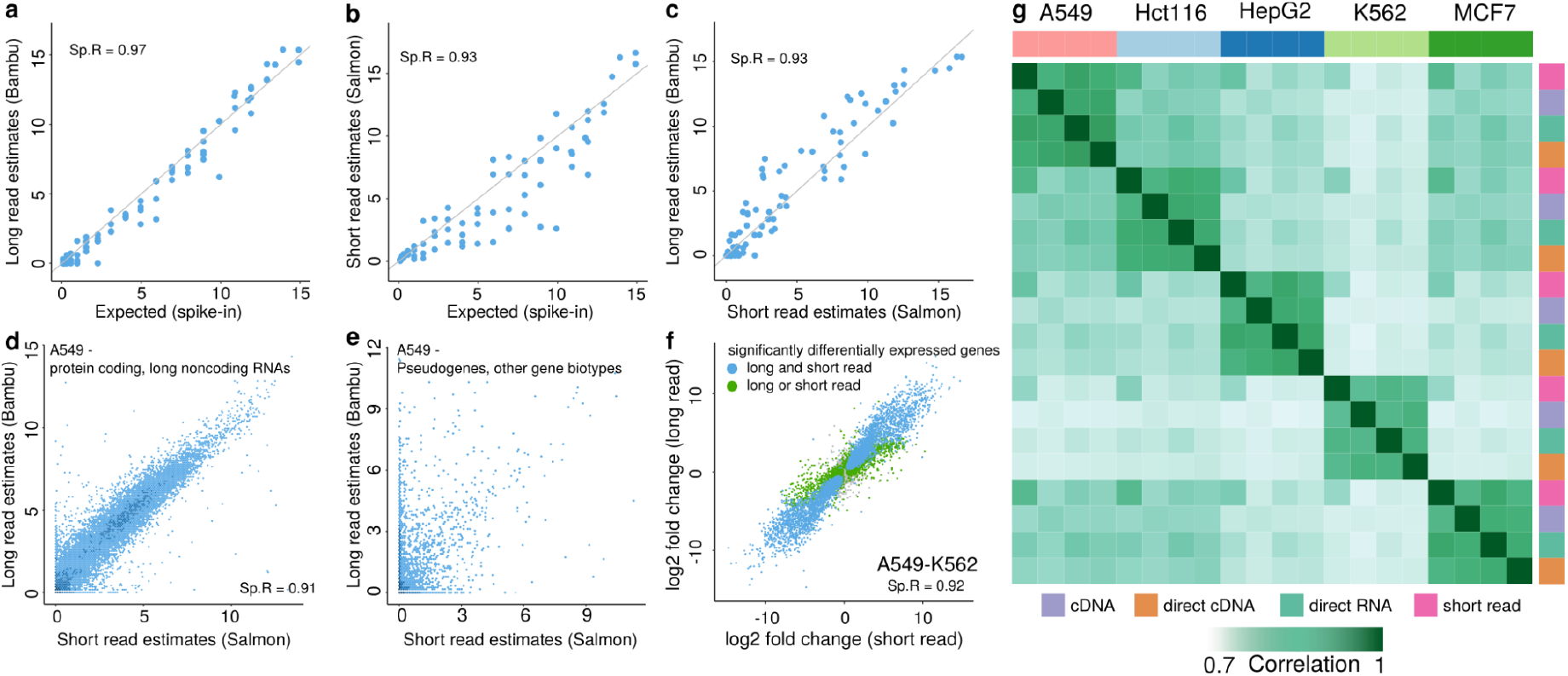
Long read RNA-Seq shows consistency in gene expression quantification with short read RNA-Seq data. **(a-b)** Scatterplots of spike-in gene expression estimates obtained from (a) long read direct cDNA RNA-Seq (using bambu) and (b) short read RNA-Seq (using Salmon with bias correction), compared against expected spike-in concentrations. **(c)** Scatterplot of spike-in gene expression estimates from long read against that from short read RNA-Seq **(d)** Scatterplot of gene expression estimates obtained from long read (using bambu) against that obtained from short read (using Salmon with bias correction) in the A549 cell lines **(e)** Scatterplot of pseudogene and other minor biotype gene expression estimates obtained from long read (using bambu) against that obtained from short read (using Salmon with bias correction) in the A549 cell lines **(f)** Scatter plot showing the log2 fold change when comparing gene expression estimates from A549 against that from K562 obtained from long read RNA-Seq against the log2 fold change from short read RNA-Seq **(g)** Heatmap showing the correlation of gene expression across the samples in the 5 cancer cell lines generated using cDNA, direct cDNA, direct RNA and short read protocols.

### (5) Long read data improves over short read data for transcript abundance estimation

While gene expression analysis is highly robust, the estimation of transcript expression abundance is more challenging as distinct transcripts from the same genes are often largely similar ^16–19^. Similar to gene expression on spike-in RNAs, we observed that long read sequencing enables more accurate quantification compared to short reads at the transcript level (R = 0.96 vs 0.94, FIgure 4a,b). However, while gene expression estimates from human cell lines were highly comparable between short and long read sequencing, transcript expression estimates show larger differences (Figures 4c,d, Supplementary Figure 3a). The highest agreement in abundance estimates is observed for the most active isoform of each gene (major isoforms, Figure 4e). However, the large difference in abundance estimates for other (minor) isoforms demonstrates that transcript level analysis with short and long read data is inconsistent (Figure 4f). Interestingly, a similar level of variation is seen when 150bp paired-end short read RNA-Seq data is compared to the identical data trimmed to 75bp single-end reads (Supplementary Figure 3b) indicating that shorter read length is the main reason for the observed variation for transcript level estimates (Supplementary Figure 3 c,d).

**Figure 4.**
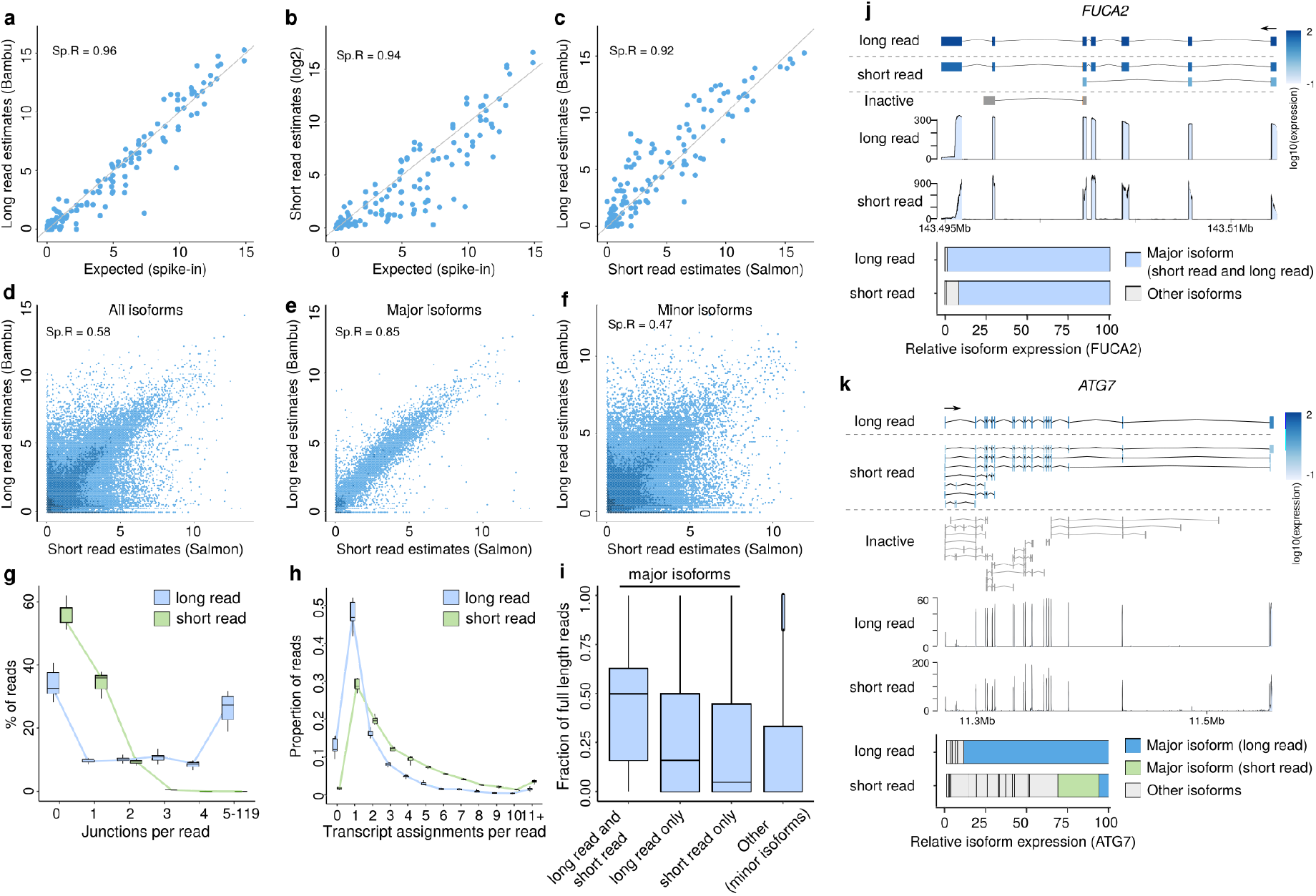
Long read RNA-Seq data improves read-to-transcript assignment and transcript abundance estimation compared to short read RNA-Seq data. **(a-c)** Scatterplots of spike-in transcript expression estimates obtained from (a) long read (using bambu) and (b) short read RNA-Seq (using Salmon with bias correction) against spike-in concentrations **(c)** Scatterplot of spike-in transcript expression estimates from long read against that from short read RNA-Seq. **(d)** Scatterplot of transcript expression estimates obtained from long read RNA-Seq (using bambu) against that obtained from short read RNA-Seq (using Salmon with bias correction) in the A549 cell lines. **(e)** Scatterplots of transcript expression estimates obtained from long read RNA-Seq (using bambu) against that obtained from short read RNA-Seq for transcripts dominantly expressed in both long read and short read in the A549 cell lines (major isoforms). **(f)** Scatterplot of transcript expression estimates obtained from long read RNA-Seq (using bambu) against that obtained from short read RNA-Seq after excluding transcripts dominantly expressed in both long read and short read in the A549 cell line. **(g)** Comparison of the number of junctions covered per read between long read and short read RNA-Seq. **(h)** Comparison of the number of transcripts uniquely assigned per read between long read and short read RNA-Seq. **(i)** Boxplot showing the fraction of reads with at least 10% full length read support for transcripts dominantly expressed in both long read and short read, transcripts only dominantly expressed in long read, transcripts only dominantly expressed in short read, and transcripts minorly expressed in both long read and short read RNA-Seq. **(j-k)** Comparison of short and long read estimates for *FUCA2* (j) and *ATG7* (k). Shown are the transcript annotations coloured by expression levels for short read and long read RNA-Seq data (top), the average read coverage (middle), and relative expression of all isoforms (bottom). Short read and long read estiamtes agree for *FUCA2*, but show major differences for more complex annotations such as *ATG7*.

Compared to short read data, Nanopore long reads cover significantly more junctions (Figure 4g), and significantly more reads can be uniquely assigned to a transcript (Figure 4h). Major isoforms identified by long read RNA-Seq show consistently higher full-length read support compared to isoforms identified by short read RNA-Seq, possibly reflecting ambiguity in read assignments related to shorter read length that could cause overestimation (Figure 4i). While genes with only few annotated transcripts show good agreement between short and long read data (Figure 4j), long reads enabled more accurate transcript abundance estimates for genes with a large number of similar transcripts (Figure 4k). Together, these data suggest that improvements in read-to-transcript assignment is one of the major benefits of using longer reads is an improvement in read-to-transcript assignment.

### (6) Full-length reads enable the analysis of individual isoform expression

The ability of long reads to unambiguously identify isoforms that are expressed enables the analysis of complex splicing events involving multiple exons. To ensure that isoforms are present in the data, we restrict this analysis to transcripts which are supported by full-length reads (see methods). Across all 5 cell lines, we observe that thousands of genes use multiple isoforms in each cell line (Figure 5a). Interestingly, the most frequent difference between alternative isoforms and the major isoform is the choice of alternative promoters (31.9%), followed by exon skipping (27.8%) and alternative last exons (18.9%), indicating that isoform usage is strongly influenced by transcriptional regulation (Figure 5b,c). Some of the most complex genes use more than 20 distinct isoforms, often involving alternative promoters, termination sites and splicing (Supplementary Figure 4a).

**Figure 5.**
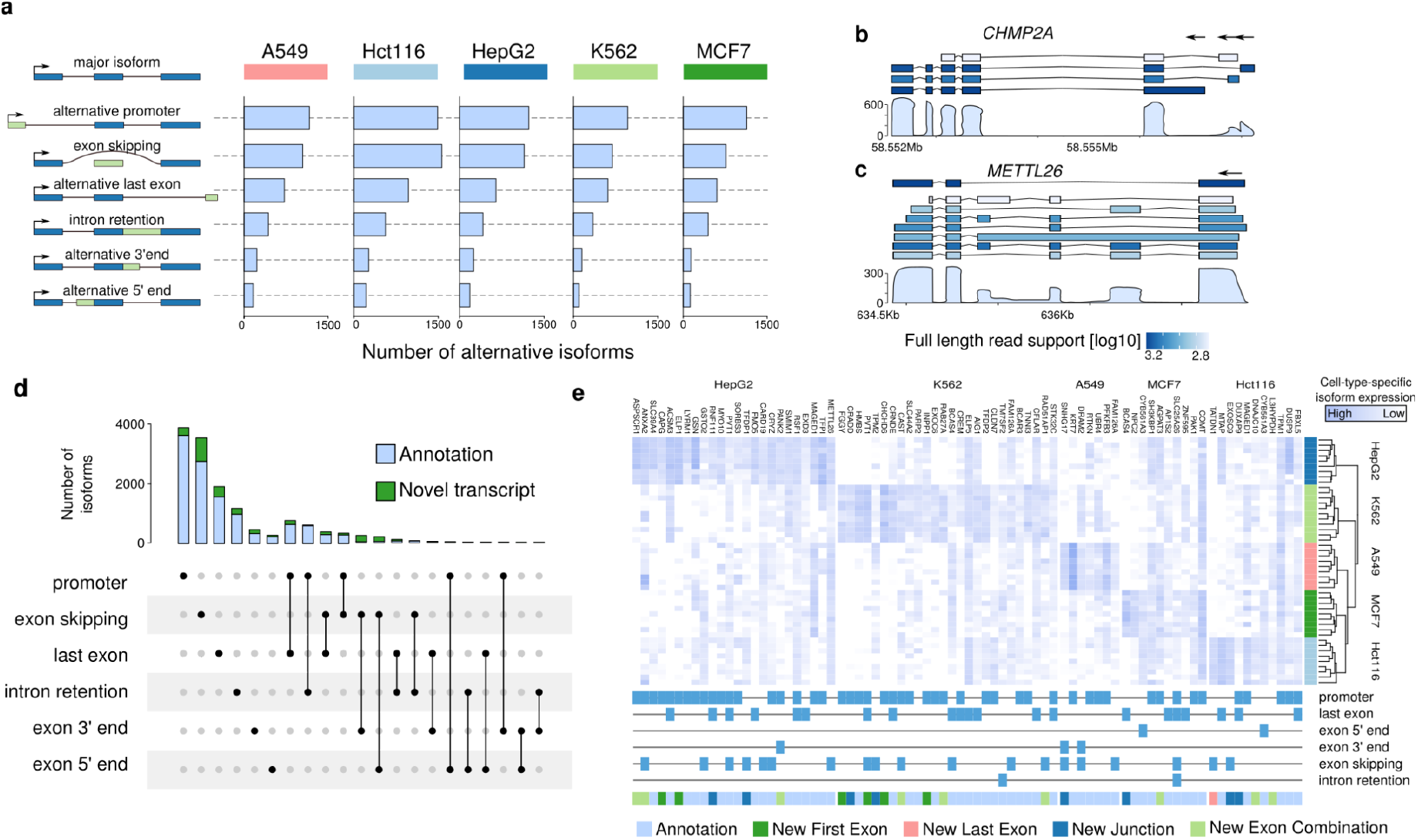
Full Length isoform analysis with long reads identifies complex transcriptional events and novel transcripts. **(a)** Barplots of different isoform switching type events in the 5 cancer cell lines. **(b)** Illustration of the annotation (top) and coverage (bottom) for *CHMP2A* with multiple promoters used. **(c)** Illustration of the annotation (top) and coverage (bottom) for *METTL26* with multiple isoform switching events happening. **(d)** Upset plot of isoform switching event combinations. Left: boxplot showing the number of isoforms for each isoform switching type across cell lines; top:number of annotated (light blue) and novel (green) isoforms for each combination. **(e)** Heatmap showing the expression levels of 231 isoforms showing significant dominant isoform switching events across the 5 cancer cell lines, the type of events associated with the isoform is indicated at the bottom. Expression is shown for the cell-type-specific isoforms.

While individual alternative splicing events and alternative promoters can be identified from short read data ^4,6,7,13^, long read RNA-Seq data enables the identification of multiple such events spanning the entire isoform. Here we find that a large fraction of isoform switching (23.9%) involves multiple events (Figure 5d). About 26.6% of alternative last exon, and about 10.6% of exon skipping events are associated with a change in promoter, illustrating how long reads provide insights into long-range associations that are not directly quantifiable with short read data.

Next we identified cell type-specific alternative isoform usage by comparing each cell line against all other cell lines from the SG-NEx data. In total we identified 231 significant isoform switching events across cell lines that involved a change in major isoform and which were supported by full length reads (Figure 5e, Supplementary Table 3, see methods). While most major isoforms are annotated, we observed several novel transcripts that were involved in isoform switching events (Figure 5e, Supplementary Figure 4b). All isoforms are supported by full-length reads, demonstrating how long read RNA-Seq enables the analysis of complex transcriptional events through the robust quantification of individual gene isoforms.

### (7) Novel genes are enriched in repetitive elements

Across all samples in the SG-NEx data we identified 4096 novel multi-exon transcript candidates, 715 of which belonged to a set of 512 unannotated genes (Supplementary Table 4). The vast majority of these new genes are of lower complexity compared to annotations, with only 13% consisting of more than 2 exons, and 12% generating multiple isoforms. Compared to annotated transcripts, we observed a significant enrichment of repetitive elements in novel transcripts (p < 0.001, Figure 6a,b). The enrichment of repetitive elements was observed for both novel transcripts from annotated genes and for novel gene candidates (Figure 6c), suggesting that one of the major advantages of long read RNA-Seq data is an improved ability to reconstruct RNAs in repeat-rich regions.

**Figure 6.**
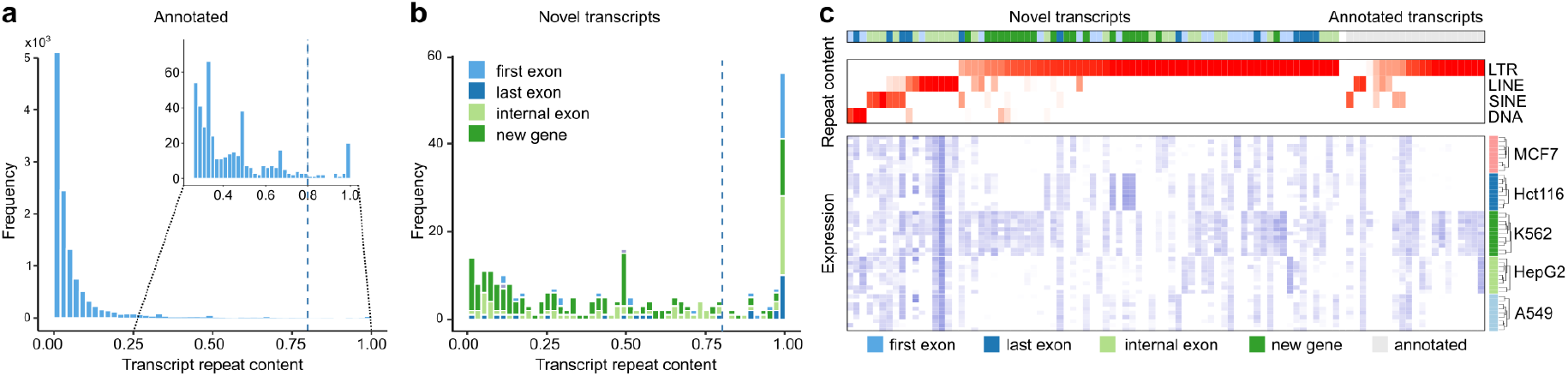
Long read RNA-Seq enables the discovery and quantification of highly repetitive genes. **(a-b)** Shown is the histogram of overlap percentage with repeat elements for annotated and novel transcripts, with new first exon, new last exon, new internal exon, and new genes shown in different colors. **(c)** Overview of the repeat family (top) and expression levels across samples in the 5 cancer cell lines (bottom) for transcripts with at least 80% overlap with major repeat elements families (LTR, LINE, SINE, and DNA).

### (8) Detection and quantification of full length fusion transcripts

The occurrence of genomic rearrangements in cancer can introduce fusion genes which are associated with clinical characteristics ^30^. Using the SG-NEx long read RNA-Seq data we searched for fusion genes in the 5 cancer cell lines. Our results indicate that long read RNA-Seq data enables the robust identification of fusion genes with 54 fusions found in the MCF7 breast cancer cell line. Across the 5 cell lines 78% of fusions discovered have been validated previously or observed in short read data (Figure 7a, Supplementary Table 5). Interestingly, we additionally find full length read support for most of the 5’ and 3’ genes, indicating that fusion chromosomes are present together with the original chromosome (Figure 7a).

**Figure 7.**
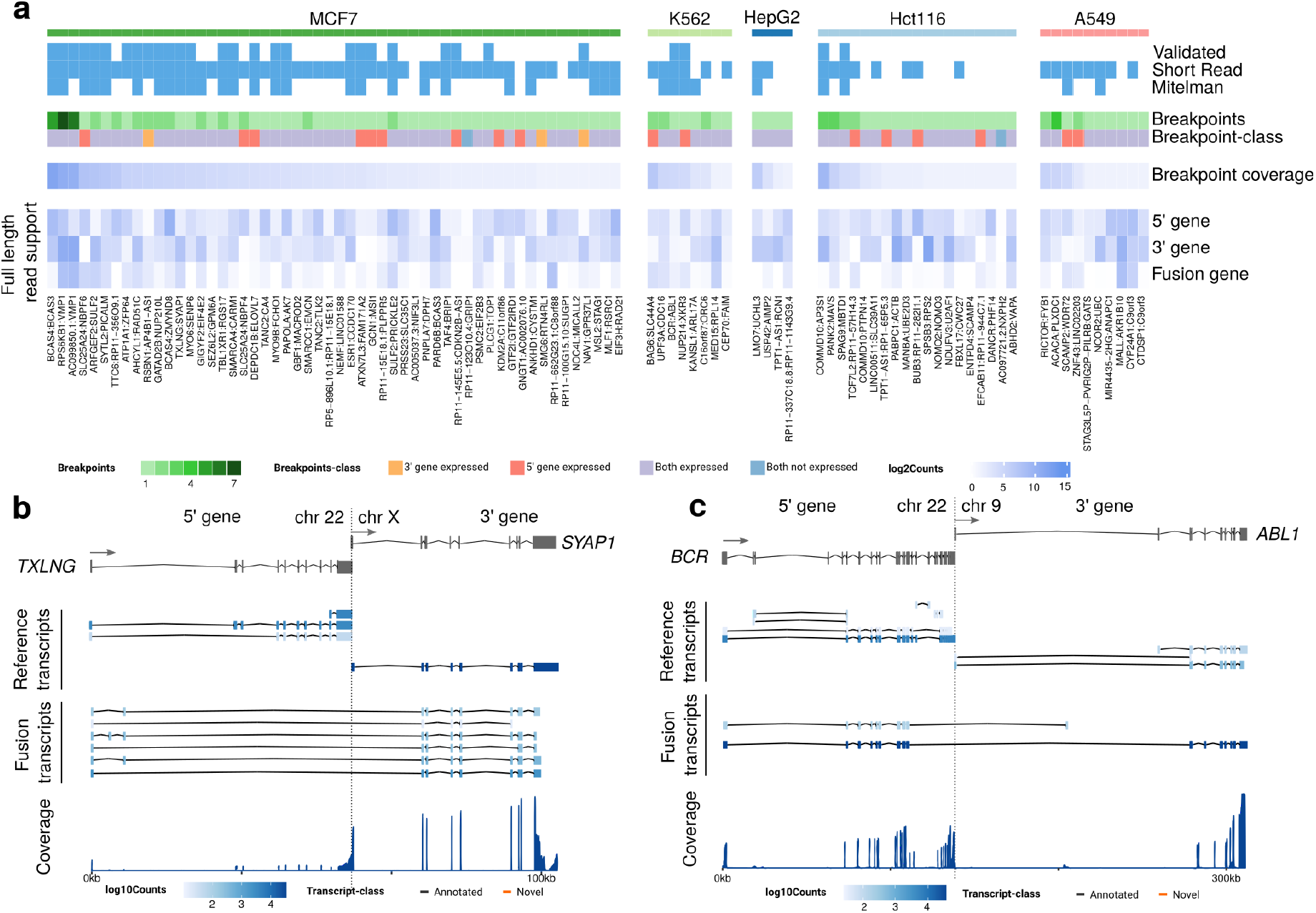
Detection and quantification of full length fusion transcripts. **(a)** Heatmap of fusion gene candidates detected using long read RNA-Seq data. Shown is the status of literature validations (top), number and class of break points (middle), and full length read support for the 5’ gene, 3’ gene and the fusion gene. **(b)** Shown are the isoform annotations for *TXLNG*, *SYAP1*, and the *TXLNG-SYAP1* fusion gene with color shading indicating the level of full length read support (top), and coverage plot in the region (bottom) in the MCF7 cell line. The *TXLNG-SYAP1* fusion gene shows alternative splicing patterns not observed in the 3’ and 5’ genes. **(c)** Shown are the isoform annotations for *BCR*, *ABL1*, and the *BCR-ABL1* fusion genes, with color shading indicating the level of full length read support (top), and read coverage plot (bottom) in the K562 cell line. Novel transcripts for 5’ and 3’ genes and similar fusion transcripts are removed for simplified visualization.

Unlike short read RNA-Seq data, which is limited to detecting the breakpoint, long read RNA-Seq data enables the reconstruction of complete fusion transcripts. This not only identifies transcripts with alternative breakpoint junctions but also enables the discovery of associated splicing events away from the breakpoint that are specific to the fusion transcripts (Figure 7b). The ability to profile complete fusion transcripts is further demonstrated for the well-described breakpoint that leads to the fusion gene *BCR-ABL1*. Using the SG-NEx data, we identify a novel *BCR* fusion transcript that uses an alternative last exon on the fusion chromosome 9, but which does not use any exon from the *ABL1* gene (Figure 7c). This novel fusion transcript is supported by reads covering all exons, illustrating how long read RNA-Seq enables the discovery and quantification of full-length fusion transcripts in cancer.

### (9) The landscape of m6A RNA modifications in the SG-NEx cell lines

The ability to directly sequence RNA using the Nanopore technology facilitates the discovery of RNA modifications that otherwise requires dedicated experimental protocols ^31^. In order to identify m6A sites across the SG-NEx cell lines we first aligned the current signals to the reference nucleotide sequences (Figure 8a) ^32^. Based on the reference-aligned segments, we then applied m6Anet to obtain a set of candidate m6A positions (Figure 8a). Across all 5 cell lines, we find 2,695 positions that are predicted to be modified in at least 1 cell line (Supplementary Table 6). As expected, m6A sites identified using direct RNA-Seq data show an enrichment at the 3’ end of transcripts and resemble the expected DRACH motif (Figure 8b, Supplementary Figure 5a). Candidate m6A sites also show a shift in the current signal that corresponds to the expected deviation due to m6A modifications, further suggesting that sites identified with direct RNA-Seq data are indeed modified by m6A (Figure 8c, Supplementary Figure 5b)^33^. Globally, we observe that m6A sites show cell type-specificity, partially due to cell type-specific expression of transcripts (Figure 8d, e, Supplementary Figure 5c). In contrast, m6a sites in transcripts expressed across all cell lines are consistently found modified (Figure 8e). Among the most heavily modified genes is the oncogene *MYC* which was shown to be regulated by m6A in cancer^34^, illustrating how direct RNA-Seq can simultaneously profile RNA expression and modifications (Figure 8f, Supplementary Figure 5d,e).

**Figure 8.**
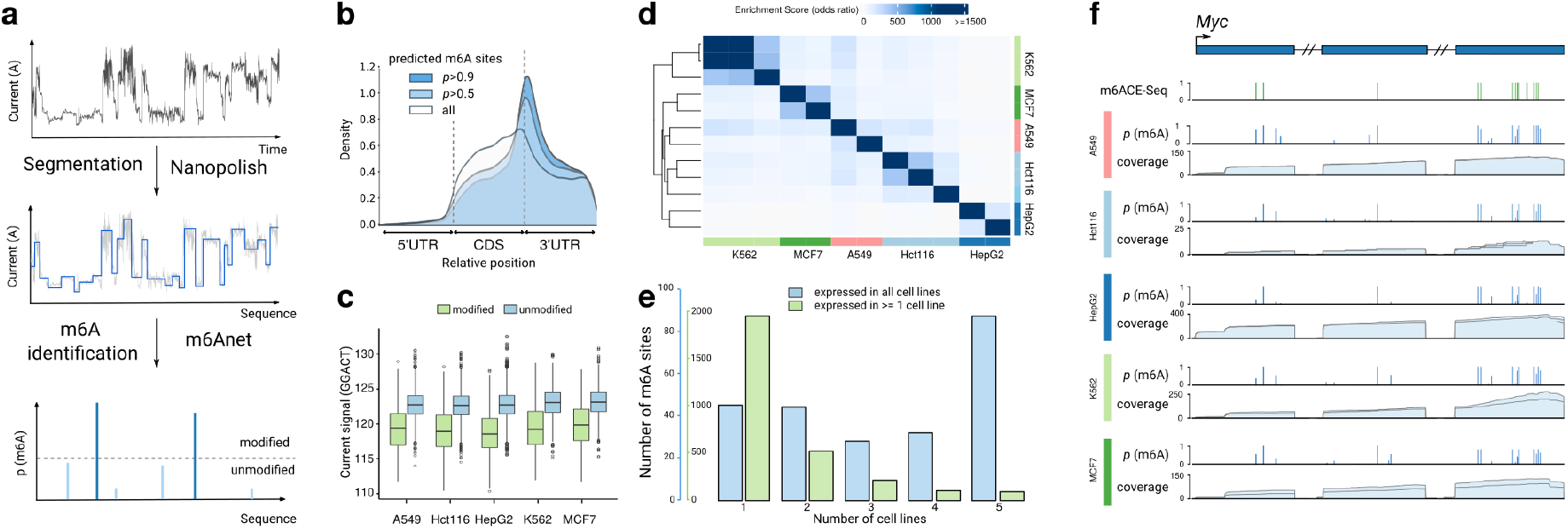
Profiling of m6A in 5 cell lines using direct RNA-Seq. **(a)** Workflow of identifying m6A positions from direct RNA-Seq data. **(b)** Metagene plot showing the frequency of m6A sites identified from direct RNA-Seq data at relative positions within the 5’UTR, CDS, and 3’UTR of a transcript. The grey line (‘all’) indicates the expected distribution based on all positions which are tested. **(c)** Boxplot showing the current signals of modified and unmodified GGACT positions across the five cell lines. **(d)** Heatmap showing the Fisher’s test odd ratio (enrichment score) across sample replicates from the five cell lines. **(e)** Barplots of shared m6A sites across cell lines, including predicted m6A sites at genes which are expressed across all cell lines (blue), and predicted m6A positions at genes which are expressed in at least 1 cell line (green). **(f)** Shown is the *MYC* gene with m6ACE-seq-detected m6A positions (green bars) and m6Anet-detected m6A probability inferred from direct RNA-Seq data (blue bars). The direct RNA sequencing coverage is shown in light blue for each cell line.

## Discussion

Here we present the results from the Singapore Nanopore Expression project, a systematic benchmark data set and comparison of 4 different RNA-Seq protocols. We find that reverse transcription to cDNA has a limited impact on transcript expression measurements, that amplification with PCR has a globally measurable impact, and that read length is the major factor influencing quantification of isoform expression. Our data suggests that long reads improve most tasks in transcriptome profiling compared to short reads while further enabling additional analyses that are only possible with long read sequencing, leading to new insights for alternative isoform regulation, repeat-derived transcripts, identification of novel isoforms associated with well-known fusion genes and profiling of m6A RNA modifications.

One of the main challenges faced by short read RNA-Seq data is a low resolution for repetitive regions ^35,36^. Highly repetitive transcripts have been shown to be expressed in embryonic development ^37^, adult tissues ^12^, and they have been associated with diseases ^38,39^, yet they are among the most difficult to study. Our data suggests that one of the main advantages of long reads is an improved resolution to identify highly repetitive transcripts, providing a more complete understanding of their role in the transcriptome. Long read data will be of particular relevance for tissues which show high expression of repeats and which are less-well annotated, such as cells from early embryos or samples from species which are known to be particularly repeat-rich ^40,41^.

Compared to short read RNA-Seq data, Nanopore RNA-Seq has a higher sequencing error rate ^24^, which affects the precision of read alignments in particular for splice junctions. Therefore, assignment of nanopore reads to transcripts requires approximate matching, providing a barrier for using tools designed for short read RNA-seq data. Error correction ^42–44^, or splice alignment correction as used in bambu^45^ Talon ^46^ and Flair ^21^ can overcome this limitation. Here we observe that the direct RNA Sequencing data, which has the highest error rate ^23,24^, generates transcript abundance estimates that are consistent with the direct cDNA data, indicating that the increased error rate can effectively be dealt with for transcript quantification.

For transcriptomic applications such as single cell profiling that rely on maximising the read count from minimal amounts of RNA, short read RNA sequencing can provide an advantage as a greater number of molecules can be detected at the same sequencing depth ^47^. However, using short reads prevents the analysis of gene isoforms at single cell resolution ^48,49^. Studies which combined long read RNA-Seq with single cell profiling have demonstrated the relevance of isoform expression ^50,51^. While the cell numbers for these studies are limited compared to short read RNA-Seq, an increased throughput can potentially enable the profiling of individual isoforms in thousands of cells with long read RNA-Seq.

One of the main advantages of short read RNA-Seq data is the availability of a large amount of public data for benchmarking and discovery ^10,52,53^. Even though the Nanopore technology is widely available and new methods are being actively developed ^54^, the lack of a comprehensive data resource still remains an obstacle towards wide-spread use of Nanopore RNA sequencing for routine profiling of the transcriptome ^54^. Here we presented the SG-NEx data set and used it to compare the different sequencing protocols. However, the data is also designed to enable comparative benchmarking of computational methods. By including multiple cell lines with multiple replicates each sequenced on all 4 protocols, the data enables the development and evaluation of methods for isoform quantification and discovery, differential expression analysis, and fusion gene detection. Furthermore, the SG-NEx data contains the raw current signal from direct RNA sequencing for all 5 cell lines each profiled with multiple replicates, providing a unique resource to develop and evaluate methods for identification of RNA modifications. Together we provide a systematic benchmark data set for short and long read cDNA and direct RNA sequencing, a comprehensive overview of transcription in human cells, and a systematic evaluation of transcriptomics protocols that highlights the benefit of long reads to studying complex transcriptional phenotypes at the resolution of individual isoforms.

## Methods

### Cell Growth

Cell growth protocols are described in Supplementary Table 1.

### RNA extraction

RNA extraction protocols are described in Supplementary Table 1.

### Library preparation

Sequencing libraries were prepared using the Nanopore direct RNA, direct cDNA, and PCR cDNA kits, and short read sequencing (Illumina). Details and deviations are described in Supplementary Table 1.

### Sequencing

Sequencing runs were performed using MinION/GridION (FLO-MIN106/106D/107), or PromethION (FLO-PRO001/002) (Oxford Nanopore Technologies) (Supplementary Table 1).

### Reference genome and annotation

We have used the Grch38 Ensembl annotations release version 91 ^1^. We used the primary assembly fasta sequence as the reference genome sequence. For transcriptome alignment we combined the coding and non-coding RNA reference fasta files, selected transcript ids that matched the reference annotations, and removed other transcripts from the transcriptome fasta file. For repeat elements, we used the matched release version of RepeatMasker sequences ^55^. For the spike-in dataset, we have used the sequin annotation ^27^. All reference files can be downloaded from https://github.com/GoekeLab/sg-nex-data.

### Basecalling and Read Alignment

Nanopore RNA-Seq data was processed following the procedure implemented in nfcore-nanoseq pipeline (nf-core/nanoseq: nf-core/nanoseq v1.0.0 - Silver Snail) ^56^. First, fast5 files are basecalled using Guppy version 3.2.10 ^57^. The resulting fastq file are then aligned using the long read aligner Minimap2 version 2.17 ^58^ with parameters “-ax splice --junc-bed” for genomic alignments using the junction bed file to correct splicing junctions, and with parameters “-ax map-ont” for transcriptomic alignments. For direct RNA-Seq runs the additional parameters “-k14” and “-uf” are used as recommended. For short reads, we performed STAR alignment with parameters “--outMultimapperOrder Random --outSAMattributes NH HI NM MD AS nM jM jI XS” to obtain the genome alignment first (which was used to calculate the junction counts).

### Transcript Discovery and Quantification

Transcript discovery and quantification was done following the nf-core/nanoseq pipeline using bambu version 1.0.2 (https://github.com/GoekeLab/bambu) with a minimum read count threshold of 2.

### Comparison of long read RNA-Seq protocols

We have compared the three long read RNA-Sequencing protocols: direct RNA, direct cDNA, and cDNA protocols in terms of sequencing depth, read length, transcript diversity, and coverage. For sequencing depth presented in Figure 2b, we have compared the total number of reads generated from each sample using different protocols. Sequencing runs with more 400,000 reads were included for the protocol comparison.

To compare the transcript diversity between protocols presented in Figure 2c, we firstly ranked the genes by the average expression within each protocol and then calculated the cumulative read frequency for genes ranked from top expressed to low expressed for each sequencing run. To summarise the transcript diversity for each protocol, we have taken the median of the cumulative read frequency across samples for each protocol.

To compare the read length, we calculated the mean read length mappable across samples within each cell line for each protocol. To evaluate the read coverage between protocols presented in Figure 2e, we have used the coverage function from GenomicAlignments ^59^ to obtain the coverage for each position along the transcript. With the default parameter, we chose to not ignore regions corresponding to D(deletion) in the CIGAR string. We then divided each transcript into 100 equal bins and aggregated the coverage read count within each bin, which was then normalized by the maximum for each transcript. We then averaged the normalized coverage ratio across samples within each protocol for each bin.

To show the distribution of read length coverage presented in Figure 2f, we calculated the coverage for each read based on the transcriptome alignment and calculated the ratio against the transcript length, and the coverage ratio will be normalized to a number over a thousands. We then calculated the mean number of transcripts for each coverage ratio ranging from 0 to 1 for each protocol and then calculated the cumulative distribution of the coverage ratios.

To compare the gene coverage presented in Figure 2h, we aggregated the read count for each read class aggregate for each gene as the expected read count if all read classes are full length with respect to the gene. We then calculated the relative length ratio of each read class where the width of each read class was divided by the maximum transcript length within the gene, and then computed a relative length-adjusted total read count for each gene as the observed read counts. By taking the ratio between the length-adjusted and original total read count, we obtained an approximate measure for gene coverage. Similarly, to compare between protocols, we averaged the gene coverage ratios across samples generated from the same protocol for each gene. We only included genes that are expressed in all protocols with an average expression level being above 30. To identify the genes that show significant differences in coverage, we have tested the proportions for each gene using a Z-test without assuming the equal variance between two protocols.

### Comparison of short read and long read data

For long read data, we used bambu on genomic alignments obtained from Minimap2 to estimate the gene and transcript expression levels. For short read data, we used Salmon version 0.11.0 with paired-end fastq files using the quasi mapping-based mode, with parameters “--validateMappings--seqBias --gcBias --posBias” to allow correcting for sequence bias, fragment-level GC bias, and the non-uniform coverage biases.

For the spike-in comparison, we have included all 6 direct cDNA sequencing runs with spike-in concentration added and all short read RNA-Sequencing runs as they all contain spike-in RNAs. Similarly, when comparing the spike-in gene and transcript estimation between using long read and short read, we have aggregated the expression levels across all direct cDNA runs with spike-in RNAs to increase the throughput for long read data, while for short read data, expression levels are averaged across all short read sequencing runs.

To compare the human chromosome gene and transcript estimation between long read and short read data, we firstly matched the annotations to include only genes with all transcripts present in both genomic annotation and transcriptomic annotations, with 1,066 (2%) genes removed. As outlined in the manuscript, we have focused on protein coding genes, antisense RNAs, linc RNAs, non coding RNAs, and macro lncRNAs, which have further removed 25,441(43.6%) genes (pseudogenes and short RNAs). After the filtering, a total of 32,861 genes are included, with 169,254 associated transcripts.

To investigate the transcript expression differences, we classified transcripts based on whether the transcript is dominant (major isoform) within a gene in long read or short read as follows: dominant in both long read and short read, dominant only in long read, dominant only in short read, and dominant in neither. We determined the dominant transcript for each gene within each cell line by ranking the average expression levels for all transcripts associated with that gene, using either long read or short read data.

We determined the number of junctions covered by each read using the GenomicAlignments package ^59^. To determine the number of reads that can be uniquely assigned, we processed short read data using bambu without transcript discovery to obtain the read class assignment. We then compared the distribution of the number of transcripts that can be assigned by each read class.

### Isoform analysis

To characterise the general isoform usage, we have compared the dominant isoform with non-dominant isoforms within each cell line and studied the usage of promoters, last exons, exon skipping, intron retention, and alternative splicing at 5’ or 3’ ends.

Isoform switching events can be generally classified into three categories: alternative promoters (alternative first exons), alternative transcription end sites usage (alternative last exons), or alternative splicing events. For alternative promoters, we compared the first exons and flagged the event as alternative promoter if the first exons are non-overlapping. Similarly, we flagged the event as alternative last exons if the last exons are non-overlapping. Alternative splicing can be of different types, alternative splicing at 3’ or 5’ ends, exon skipping event, or intron retention event. An exon skipping event is identified by checking if any of the exon ranges of the reference dominant isoform is contained within the junction ranges of the alternative dominant isoforms. Similarly, an intron retention event is identified by checking if any of the junction ranges of the reference dominant isoform are contained within the exon ranges of the alternative dominant isoforms. Alternative 5’ or 3’ splicing is flagged if exons spliced differently at 3’ or 5’ ends.

### Differential transcript usage

For differential gene expression analysis, we used DESeq2^60^. For differential transcript/promoter/transcription end site usage, we used DEXSeq^61^. In the filtering step, we used DRIMSeq ^62^ to filter genes with no more than 10 reads aggregated across at least 12 samples, which is the largest sample size for one of the cell line group, and isoforms with no more than 10 reads across at least 9 samples, the smallest sample size for one of the cell line group. The minimum feature ratio to gene is default to be 0.1. Although the above filtering criterion could identify important differentially used isoforms across cell lines, it also identifies negative events where genes were only expressed in non-reference cell lines as the non-reference cell lines are of larger sample size than the required sample size. To fix this issue, we implemented another filtering criterion before the above filtering criterion, where we removed genes without any expression in the reference cell line. After identifying the genes and transcripts that contribute to the isoform switching events, we checked whether the contributing transcripts are dominant isoforms of the reference cell line by finding the most highly expressed isoform within each gene using the average expression levels to decide whether it is a dominant isoform switching event.

For dominant isoform switching events, we compared the isoform from the reference cell line (reference dominant isoform) and the dominant isoforms from other cell lines (alternative dominant isoform). We then determined the isoform switching types.

### Repetitive gene analysis

To study the overlap between transcripts and repeat elements, we looked at the extended annotation set obtained using the SG-NEx cell line samples. This analysis was performed at the exon level, distinguishing between annotated or novel based on whether the exon is overlapping with annotations. We then calculated the overlapping percentage between each of the exons and the repeat elements. For each transcript, we averaged the overlapping percentages for each repeat by the exon annotation status, which will then be summed up to provide the total overlapping percentage for the transcript. We then focused on transcripts with at least 80% overlap with repeat elements.

### Fusion gene analysis

#### Fusion calling method

For each cell line in the SG-NEx dataset, reads from replicates were combined and analysed with the JAFFAL pipeline of JAFFA (version 2.0, https://github.com/Oshlack/JAFFA) ^63^. Briefly, JAFFAL detects fusions in long read data by aligning reads to a reference transcriptome (Gencode version 22) using minimap2. Reads aligning to multiple genes are flagged as candidate fusion reads and aligned to the human reference genome, hg38, using minimap2. Fusions with breakpoints within 300kbp of each other and the 5’ and 3’ genes in transcriptional order, are consistent with run-through transcription and were removed. Fusions involving genes on the mitochondrial chromosome or with low read support (<3) were also removed. Four fusions found across multiple cell lines were likely to be false and hence only those unique to a cell line were retained. Fusions previously validated in the same cell lines were identified from a literature search (Supplementary Table) ^64–73^. Fusions were identified in the Illumina 150bp paired-end data by running JAFFA’s Direct pipeline on each replicate of each core cell line.

#### Fusion alignment

With the identified fusion gene candidates, we first created a fusion annotation set that combined the sequences of the two fusion genes. We then aligned reads overlapping with any of the candidate genes in the fusion gene set to the fusion gene sequences. We then run bambu on those fusion alignments to discover fusion transcripts and quantify the fusion transcript abundance. To quantify 5’ genes, we aggregated full length read support for all transcripts that fell only within the 5’ gene range. Similarly, we aggregated the full length support for the 3’ gene. Full length read counts for transcripts that overlapped with the fusion gene range but not contained only within 5’ or 3’ gene ranges were aggregated as the full length read support for the fusion gene.

#### m6A modification analysis

For each sample in the SG-NEx dataset, signal events were aligned to a reference sequence using nanopolish ^32^. The reference-aligned signal events were analysed with m6Anet (https://github.com/GoekeLab/m6anet), which predicts the probability of DRACH motifs being modified. The following analyses were done on DRACH positions with support by ≥ 100 reads (Supplementary Table 6). The relative position of the predicted m6A sites (all, probability > 0.5, or probability > 0.9) and the density at each relative position were calculated with Python packages seaborn and matplotlib (Figure 8b). For the remaining analysis, we defined DRACH motifs with an m6Anet-predicted m6A probability ≥ 0.9 as modified (m6A sites) and otherwise unmodified. The nucleotide content of modified and unmodified m6A sites was visualized with the ggseqlogo R package ^74^ (Supplementary Figure 5a). The mean current signals of the three most common DRACH motifs (GGACT, GAACT, and ACACT) was visualised using boxplots that show the median, 25th and 75th percentiles, and the smallest and largest values within 1.5 times interquartile range below the 25th percentile and above the 75th percentile, respectively (Figure 8c, Supplementary Figure 5b). The similarity across samples was examined using the ComplexHeatmap R package ^75^ with default parameters to cluster the odd ratio from the Fisher’s test (Figure 8d) and Pearson correlation of all samples (Supplementary Figure 5c). We first averaged the predicted probability of replicates with at least 100 reads from the same cell line, then we counted the number of shared m6A sites across cell lines with two scenarios: (1) including only positions expressed across all cell lines (blue) and (2) including positions that expressed in more than one cell line (green) (Figure 8e). In order to rank genes by m6A modification status, we counted the number of m6A sites of each gene in all samples (Supplementary Figure 5d). The Myc gene’s m6ACE-detected m6A sites, cell-line-averaged m6Anet-predicted m6A probability, and cell-line-averaged coverage were plotted using the R packages sushi and ggplot2 (Figure 8f). All m6A positions of the top 50 genes across all samples were plotted as a heatmap with a color gradient indicating the predicted m6A probability using the gplots R package with default parameters for clustering the m6A positions and the cell lines (Supplementary Figure 5e).

## Data availability

The sequencing data generated in this study is available online: https://github.com/GoekeLab/sg-nex-data

## Software Availability

The nextflow pipeline nf-core/nanoseq is a streamlined, community curated pipeline for Nanopore Sequencing data processing and analysis: https://github.com/nf-core/nanoseq

## Acknowledgments

This work is funded by the Agency for Science, Technology and Research (A∗STAR), Singapore. We would like to acknowledge Shyam Prabhakar and Niranjan Nagarajan for help with data generation.

## Supplementary Figures

**Supplementary Figure 1:**
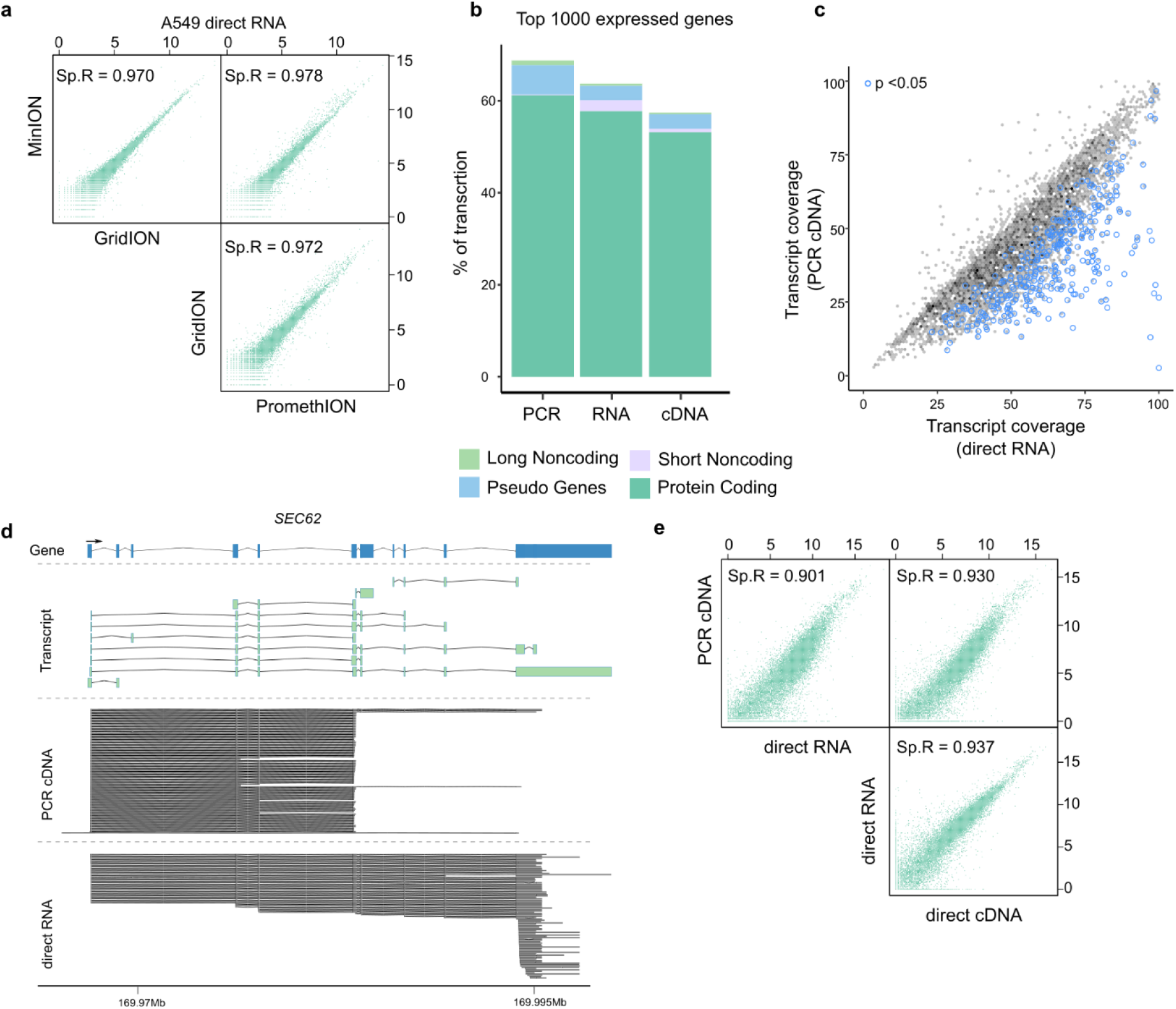
Comparison of Nanopore RNA-Seq protocols and platforms. **(a)** Shown is the comparison of gene expression generated on MinION, GridION, and PromethION platforms for A549 direct RNA samples **(b)** Shown is the biotypes for the top 1000 expressed genes generated using RNA (direct RNA), cDNA (direct cDNA) and PCR (cDNA) protocols **(c)** Shown is the transcript coverage of reads generated using the direct RNA and the PCR cDNA protocol. Blue points indicate genes with significantly reduced coverage in the PCR cDNA protocol **(d)** Comparison of reads generated using PCR cDNA and direct RNA protocols for the *SEC62* gene **(e)** Shown is the comparison of average gene expression across samples generated using PCR cDNA, direct RNA and direct cDNA protocols

**Supplementary Figure 2:**
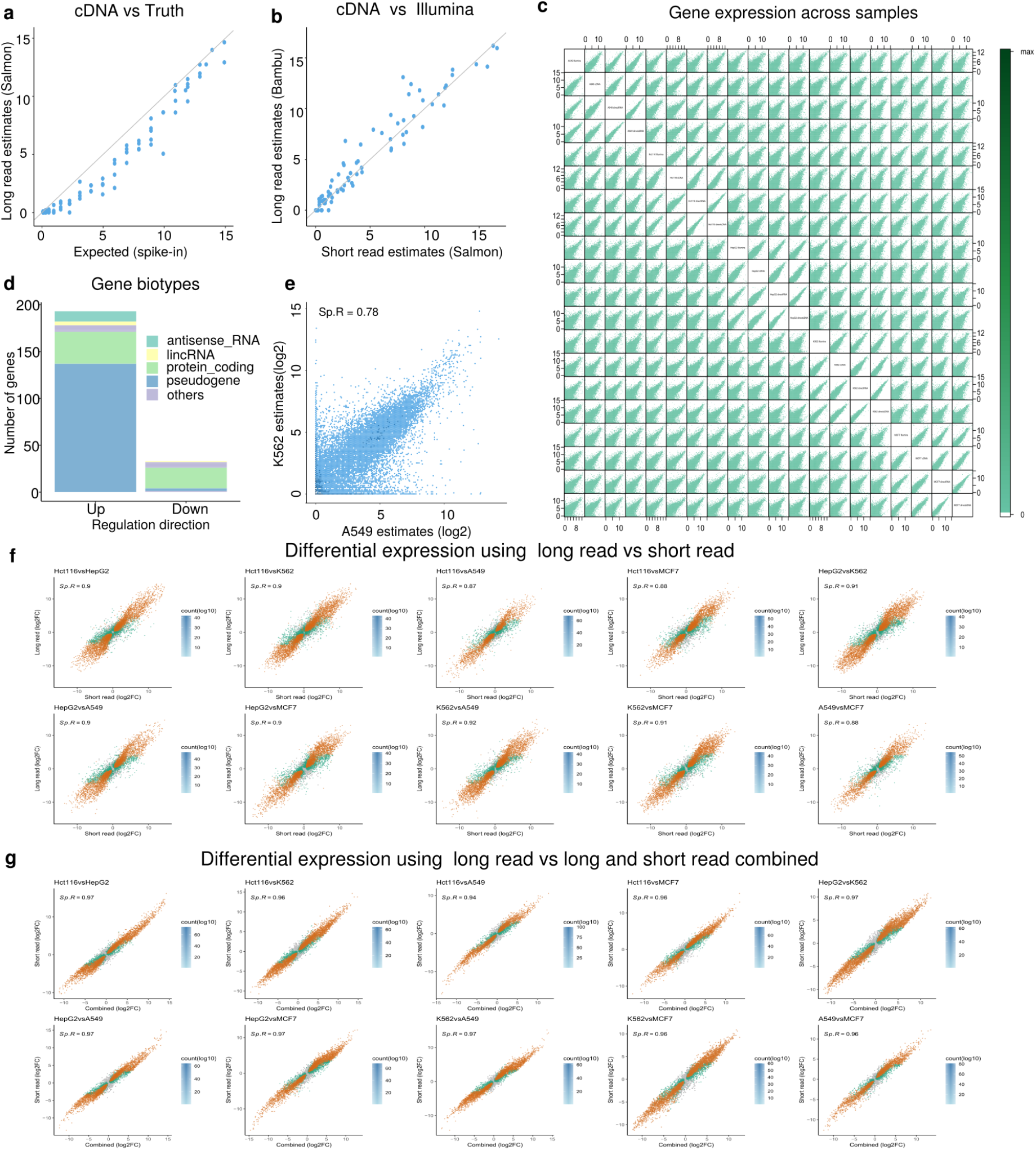
Long read RNA-Seq shows consistency in gene expression quantification compared to short read RNA-Seq. **(a)** Scatterplots of spike-in gene expression estimates obtained from long read RNA-Seq (using Salmon) compared against expected spike-in concentrations **(b)** Scatterplot of spike-in gene expression estimates from long read cDNA samples against that from short read RNA-Seq **(c)** Scatter matrix showing the gene expression across the samples in the 5 cell lines generated using cDNA, direct cDNA, direct RNA and short read protocols **(d)** Shown is the barplot of gene biotypes for genes that show significant difference in expression levels comparing long read to short read, Up meaning up regulated in long read. **(e)** Scatterplot of transcript expression estimated from 2 different cell lines **(f)** Scatter plots showing the log2 fold change when comparing gene expression estimates from two different cell lines using long read RNA-Seq against the log2 fold change using short read RNA-Seq **(g)** Scatter plots showing the log2 fold change when comparing gene expression estimates from two different cell lines using long read RNA-Seq against the log2 fold change using combined long and short read RNA-Seq

**Supplementary Figure 3:**
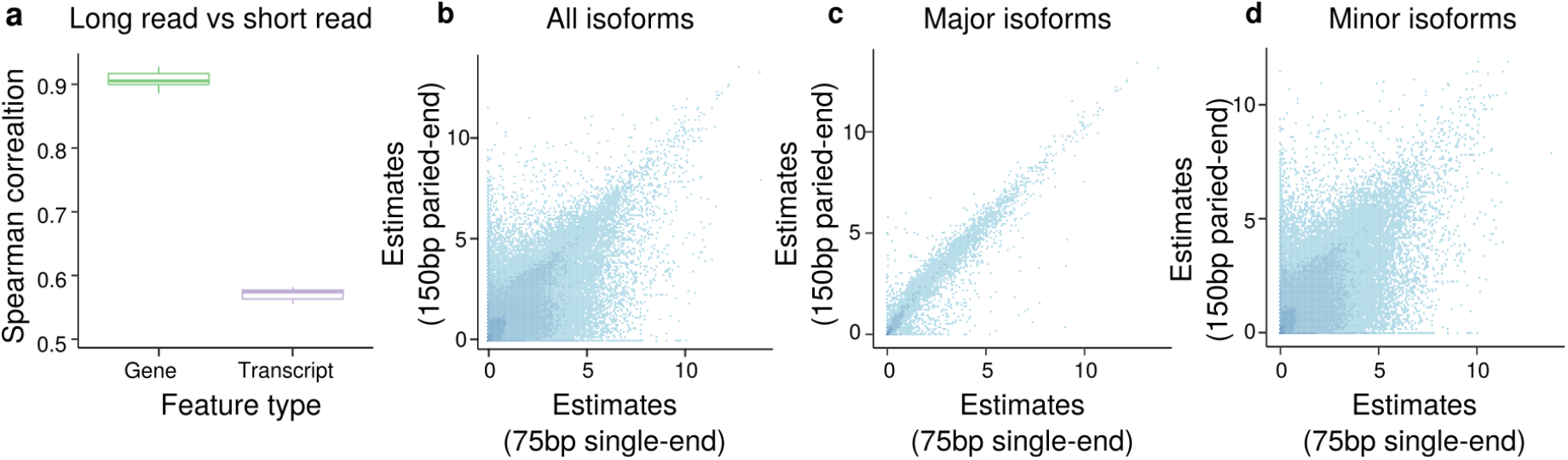
Long read RNA-Seq data improves read-to-transcript assignment and transcript abundance estimation compared to short read RNA-Seq data. **(a)** Shown is the boxplot of correlation between gene or transcript expression obtained from long read (using bambu) and that obtained from short read (using Salmon with bias correction) across cell lines **(b)** Scatterplot of transcript expression estimates obtained from 150bp paired-end short read RNA-Seq (using Salmon with bias correction) against that obtained from 75bp single-end short read RNA-Seq (using Salmon with bias correction) in the A549 cell lines. **(c)** Scatterplot of transcript expression estimates obtained from 150bp paired-end short read RNA-Seq (using Salmon with bias correction) against that obtained from 75bp single-end short read RNA-Seq (using Salmon with bias correction) for transcripts dominantly expressed in both long read and short read in the A549 cell lines (major isoforms). **(d)** Scatterplot of transcript expression estimates obtained from 150bp paired-end short read RNA-Seq (using Salmon with bias correction) against that obtained from 75bp single-end short read RNA-Seq (using Salmon with bias correction) after excluding transcripts dominantly expressed in both long read and short read in the A549 cell line.

**Supplementary Figure 4:**
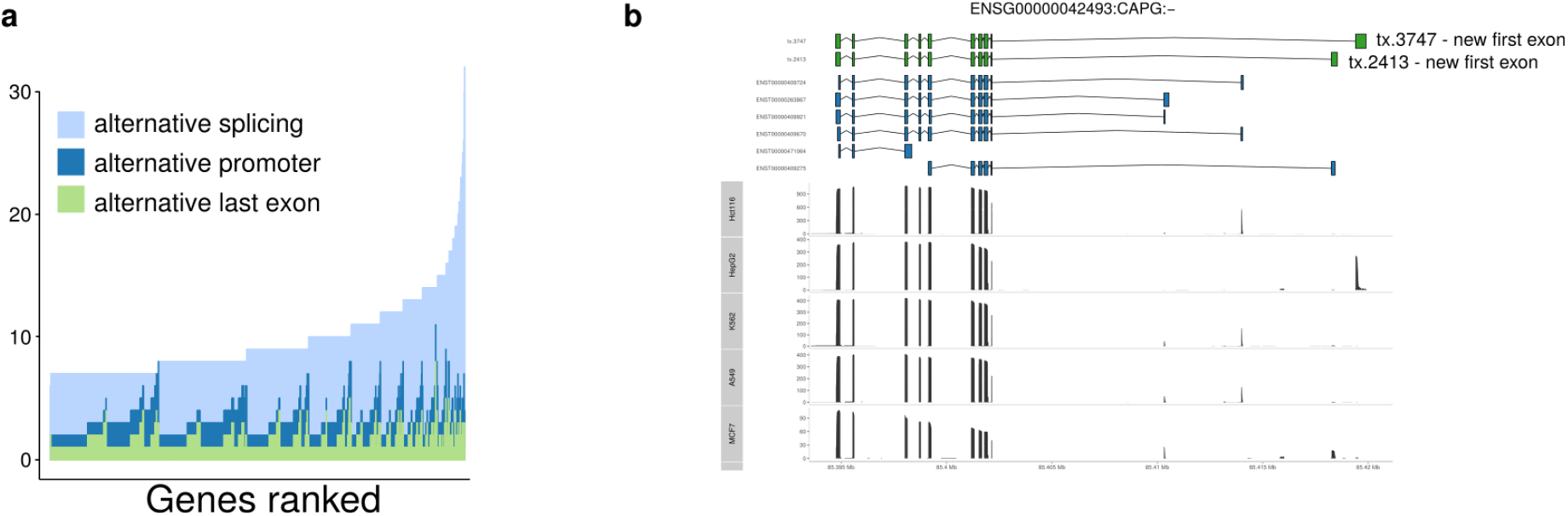
Full Length isoform analysis with long reads identifies complex transcriptional events and novel transcripts. **(a)** Barplots of genes ranked by number of isoforms expressed, with number of promoters colored in dark blue and number of last exons colored in green.) **(b)** Example of a gene (CAPG) that shows alternative isoform expression driven by alternative promoters, including 2 novel transcripts (green). Annotations are shown in blue, the read coverage is shown black.

**Supplementary Figure 5.**
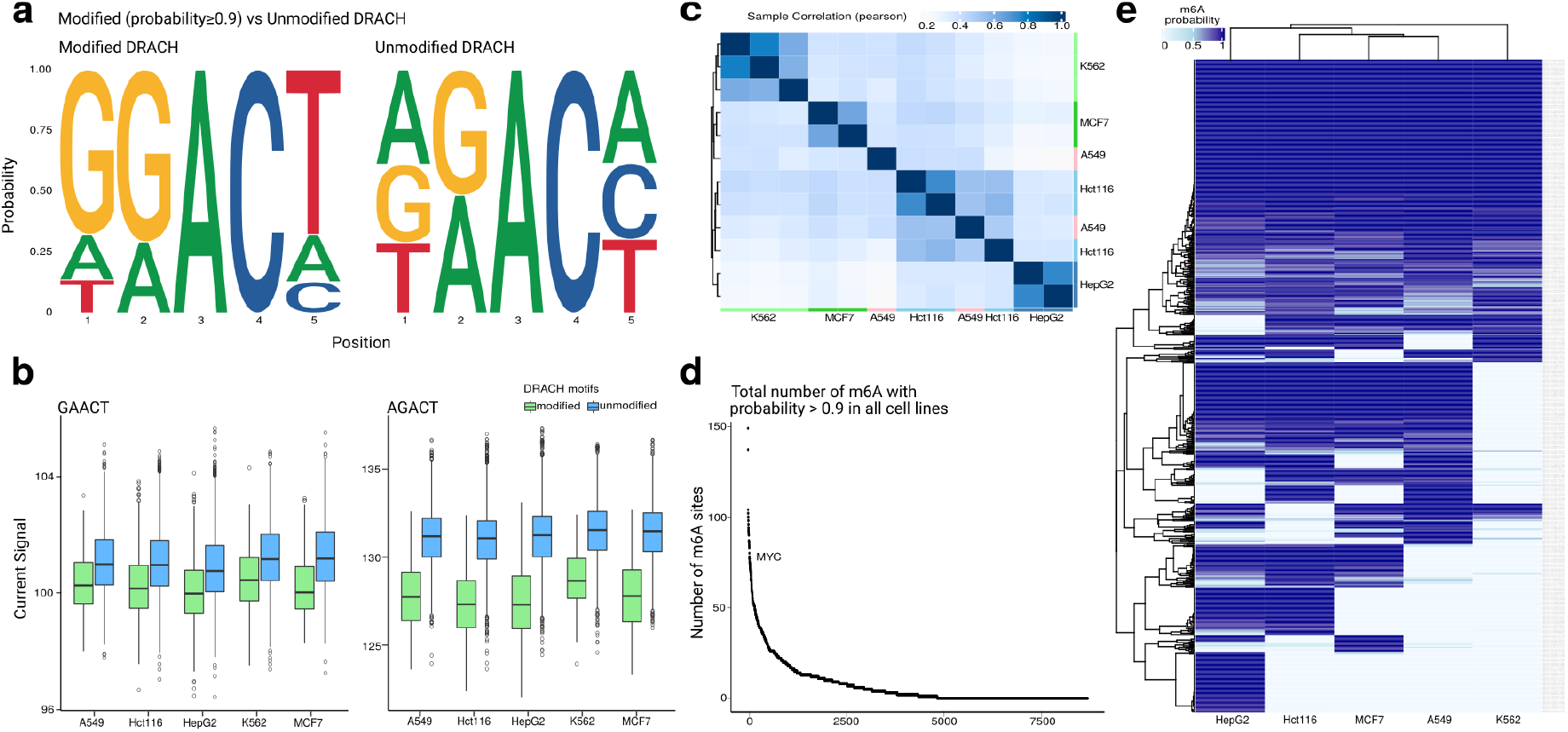
Profiling of m6A in 5 cell lines using direct RNA-Seq. **(a)** Nucleotide frequency at the five positions of all k-mers from candidate m6A positions identified by m6Anet (left) and from positions which were not identified to be modified (right). Only DRACH motifs were scanned for m6A. **(b)** Boxplot showing the current signals of modified and unmodified GAACT and AGACT positions across the five cell lines. **(c)** Heatmap showing the Pearson correlation across sample replicates from the five cell lines. **(d)** Genes ranked by the total number of m6A sites across cell lines. *MYC* ranks 38. **(e)** Heatmap of m6Anet-predicted probability of m6A sites found in top 50 genes with the highest total number of m6A sites across cell lines.

## Supplementary tables

Supplementary Table 1: Library preparation, RNA extraction and sequencing platforms across samples

Supplementary Table 2: Genes show significant lower coverage in PCR cDNA

Supplementary Table 3: Expression levels of isoform switching events

Supplementary Table 4: Expression levels of novel transcripts

Supplementary Table 5: Expression levels of fusion genes

Supplementary Table 6: predicted m6A sites in the SG-NEx data

